# Loss of Recurrent Excitation Disrupts Sleep Slow Oscillations in the Aging Brain

**DOI:** 10.64898/2026.03.16.712170

**Authors:** M. Gabriela Navas Zuloaga, Shaun M. Purcell, Maxim Bazhenov

## Abstract

Sleep-dependent memory consolidation relies on slow oscillations (SOs) that coordinate thalamocortical-hippocampal dynamics during slow-wave sleep (SWS). Aging disrupts SO properties, reducing SO amplitude, density, and slope, yet the circuit-level mechanisms linking structural brain changes to these disruptions remain poorly understood. Here we present a multi-scale, whole-brain thalamocortical network model incorporating biologically grounded human connectivity derived from diffusion MRI tractography, comprising over 10,000 cortical columns per hemisphere with spiking pyramidal and inhibitory neurons and an anatomically differentiated thalamic network. Simulating progressive synaptic loss, we find that selective degradation of recurrent excitatory connectivity, but not excitatory-inhibitory projections, reproduces empirically observed age-related SO changes. Increased SO duration was driven primarily by prolonged Down states, while Up state duration and spike density were reduced, suggesting a possible mechanism for impaired memory consolidation. These results suggest that aging selectively disrupts the temporal structure of SWS critical for interference-free memory consolidation, providing mechanistic insight into cognitive decline in the aging brain.

Supported by: NIH (grants 1R01MH125557, 1RF1NS132913, 1R01AG099626 to MB)

## 1. Introduction

Sleep is essential for long-term memory consolidation and cognitive integrity. Slow-wave sleep (SWS), the dominant brain state of non-rapid eye movement (NREM) sleep, is characterized by large-amplitude, low-frequency activity in the electroencephalogram (EEG). Its defining electrophysiological hallmark, the slow oscillation (SO; ∼0.5–1 Hz), reflects alternating depolarized (“up”) and hyperpolarized (“down”) states in the thalamocortical network (Steriade et al., 1993).

These oscillations temporally coordinate thalamic spindles and hippocampal sharp-wave ripples, providing a mechanistic framework for systems memory consolidation (Diekelmann & Born, 2010; Klinzing et al., 2019). Experimental enhancement of slow oscillations in young adults improves memory performance (Marshall et al., 2006), underscoring their causal role in sleep-dependent plasticity.

Aging alters both brain structure and sleep physiology. After the age of 40, total brain volume declines at an estimated rate of ∼5% per decade (Markov et al., 2022), accompanied by cortical thinning and progressive white matter degradation (Dubé et al., 2015). In parallel, older adults exhibit marked changes in SWS architecture and SO dynamics, including reduced amplitude, density, and slope, as well as prolonged Up and Down states (Carrier et al., 2011; Djonlagic et al., 2021). Age-related disruptions in slow-wave activity have been linked to impaired memory consolidation and cognitive decline (Djonlagic et al., 2021; Helfrich et al., 2018; Mander et al., 2013). Moreover, structural atrophy of medial prefrontal cortex predicts diminished SO generation and memory retention in older adults (Mander et al., 2013), suggesting a direct link between anatomical degradation and deficits in sleep-dependent cognitive deficits.

Despite converging evidence that structural aging and sleep disruption are intertwined, the circuit-level mechanisms connecting large-scale anatomical changes to altered SO dynamics remain unclear. Empirical studies establish correlations between cortical thinning, white matter integrity, and sleep oscillatory properties, but cannot disentangle causality or identify which synaptic pathways are most critical. Biophysically grounded computational models offer a principled framework to bridge this gap by linking structural connectivity to emergent thalamocortical dynamics. Prior modeling work has simulated aging as reduced cortical synaptic connectivity, reproducing aspects of age-related SO degradation and associated memory impairment (Wei et al., 2023). However, these approaches have been limited in spatial scale and anatomical realism, constraining their ability to capture how distributed structural degeneration shapes global sleep dynamics.

Building on this foundation, we developed a multi-scale, whole-brain thalamocortical network model incorporating biologically grounded human connectivity. The model comprises 10,242 cortical columns per hemisphere, organized into six layers with spiking pyramidal (PY) and inhibitory (IN) neurons, and an anatomically differentiated thalamic module including thalamocortical (TC) and reticular (RE) neurons within core and matrix systems. Long-range connectivity and distance-dependent delays are derived from diffusion MRI tractography, enabling large-scale structural constraints on emergent dynamics. The model reproduces both wake-like activity and spontaneous slow oscillations.

To investigate how structural aging shapes SO dynamics, we simulated progressive synaptic loss and compared resulting oscillatory properties with human EEG data across age cohorts. We show that selective degradation of recurrent excitatory (PY–PY) connectivity — but not excitatory–inhibitory (PY–IN) projections — best recapitulates the empirically observed age-related reductions in SO amplitude, density, and slope, as well as increased oscillation duration. These findings identify preferential loss of recurrent excitation, and a consequent shift in excitation–inhibition balance, as a plausible mechanistic substrate for age-related sleep disruption. Notably, the increase in overall SO duration was driven primarily by prolonged Down states, while Up state duration and spike density were reduced. These findings suggest that aging selectively disrupts the temporal structure of SWS in ways that may impair interference-free memory consolidation, offering new mechanistic insight into cognitive deficits in the aging brain.

## 2. Results

### 2.1. Realistic whole-brain cortical connectivity

We first summarize the architecture of the multi-scale thalamocortical model (Figure 1; see Methods). The cortical sheet comprises 10,242 columns uniformly distributed across one hemisphere, corresponding to vertices of the ico5 surface reconstruction described in (Rosen et al., 2019). Each column contains six cortical layers (L2–L6, Figure 1a) populated by computationally efficient map-based (Komarov et al., 2018) spiking pyramidal (PY) and inhibitory (IN) neurons. Intra-columnar connectivity follows the canonical cortical circuit (Figure 1b).

**Figure 1.**
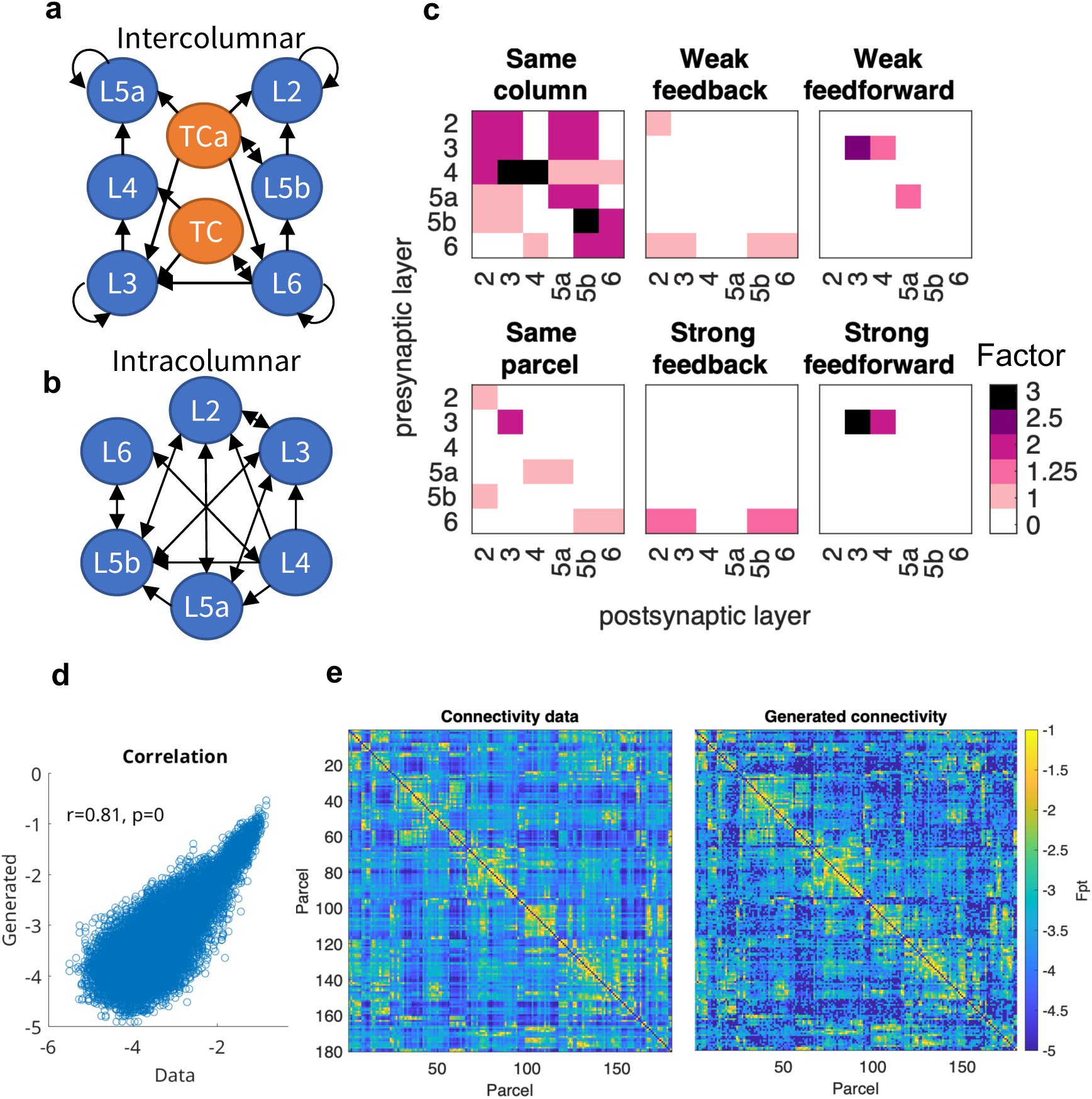
a) Connectivity within columns follows the canonical circuit. Arrows indicate directed connections between layers. Neurons are never connected to themselves. b) Connectivity between different cortical columns and core (TC) or matrix (TCa) thalamocortical cells. c) Excitatory corticocortical connections belong to one of 6 classes based on the myelination-based hierarchical index of the pre- and post-synaptic neurons: within the same column, within the same parcel, weakly or strongly feedforward (from lower to higher hierarchical index) and weakly or strongly feedback (from higher to lower hierarchical index). Based on experimental reports on connectivity (Markov et al., 2013; Rockland, 2019) weights are scaled according to the connection type by the factor shown in the matrices. d) Correlation between generated and original connectivity between parcels (Pearson’s correlation r = 0.81, p < 0.0001). e) Parcel-wise connectivity from (Rosen & Halgren, 2021) (left) and generated model connectivity (right), which retains the parcel-wise structure from data. Adapted from (Marsh et al., 2024).

Long-range corticocortical connectivity is derived from diffusion MRI tractography of the Human Connectome Project young adult dataset (Van Essen et al., 2013), organized according to the 180 parcels in the HCP multimodal atlas (Glasser et al., 2016). Parcel-level connectivity statistics from Rosen C Halgren (2021) were mapped onto column-wise distances to generate probabilistic intercolumnar projections. The originating and terminating layers of individual connections are assigned according to the parcels’ relative myelination-based hierarchical positions (Figure 1c). The resulting network preserves the parcel-wise structure of empirical connectivity and exhibits an approximately exponential decay of connection probability with distance, consistent with anatomical observations. Conduction delays scale proportionally with connection length and follow a similar distribution.

The thalamus includes thalamocortical (TC) and reticular (RE) neurons organized into core and matrix systems, reproducing known projection patterns within cortico-thalamo-cortical loops (Bazhenov et al., 1998a, 1998b; Bonjean et al., 2011). Neuromodulatory influences of acetylcholine, norepinephrine, and GABA were implemented following (Krishnan et al., 2016) to enable transitions between wake and slow-wave sleep (SWS).

To validate large-scale connectivity, we compared the generated connectivity matrix with the empirical parcel-wise matrix (Figure 1d-e). The model connectivity showed significant correlation with empirical data (Pearson’s correlation r = 0.81, p < 0.0001), indicating preservation of biologically meaningful topological structure.

### 2.2. Variable synchrony in slow oscillations

Under wake-state parameterization, the network exhibited asynchronous irregular activity with mean firing rates of approximately 12 Hz (Figure 2a). Single-neuron membrane potentials showed irregular spiking with a unimodal voltage distribution centered near resting potential.

**Figure 2.**
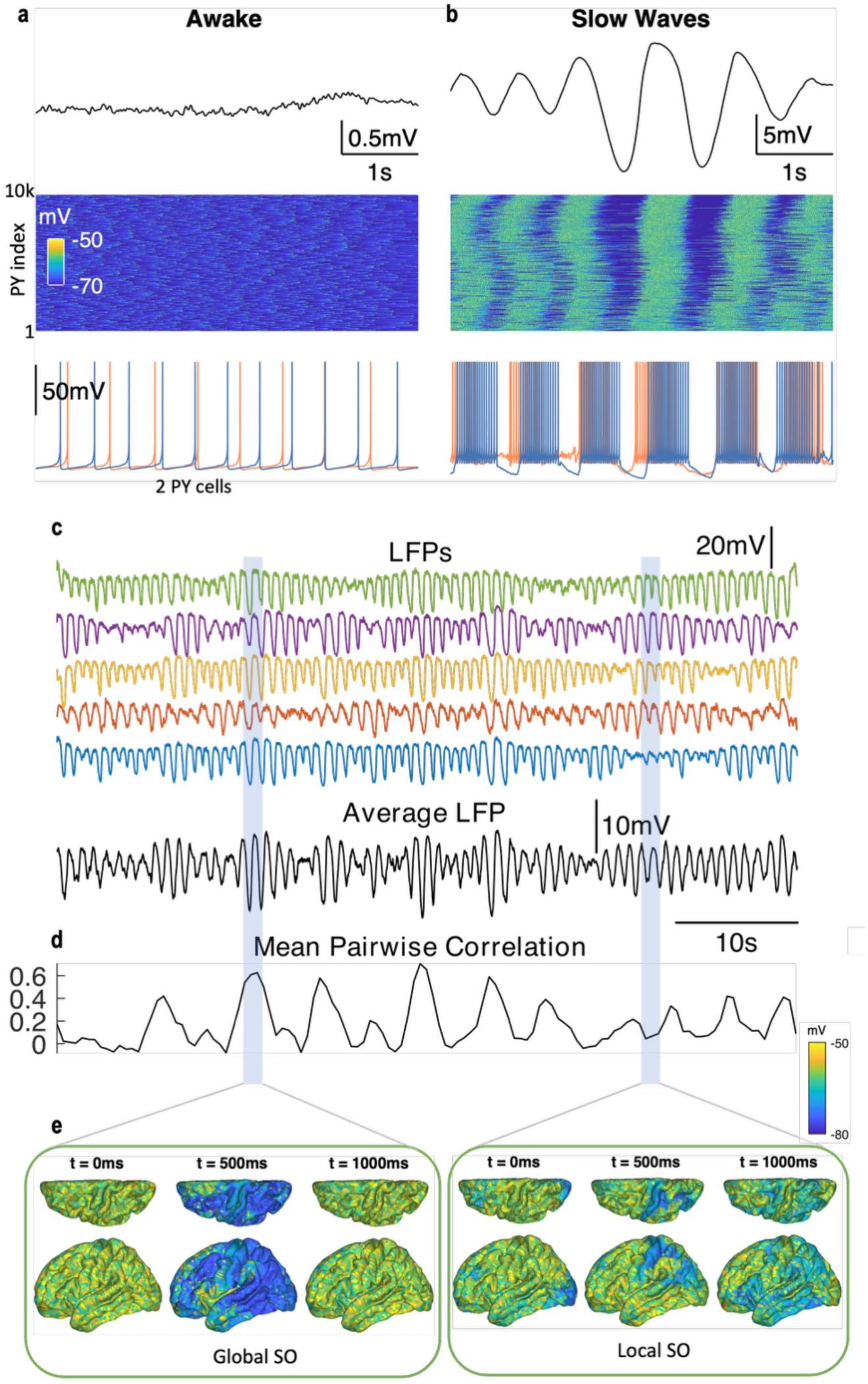
a) Model dynamics in the Awake state, with LFP (top), voltage heatmap (middle), and two pyramidal cell traces (bottom). b) Model Slow-wave Sleep (SWS) dynamics, panels as a). c) SWS activity for 5 different 5mm regions (top) and average voltage signal (bottom). d) Mean pariwise correlation between 10 LFPs. e) Snapshots of voltage for a Global SO (left) with highly correlated activity, and a Local SO (right) with low correlation between local regions.

Transition to SWS activity from the wake state involved several cellular and neurochemical parameters adjusted according to the methodology in (Krishnan et al., 2016), namely i) increased cortical AMPA, GABA, and potassium leak currents, ii) increased thalamic GABA, and TC cell leak currents, and iii) decreased RE cell leak currents. Transitioning to SWS produced robust slow oscillations (Figure 2b).

During SWS, LFPs alternated between periods of widespread synchrony and relative desynchronization, giving rise to both global and local SOs (Figure 2c). Global events were marked by long, spatially extensive down states (Figure 2e). Sliding-window correlations revealed substantial fluctuations in inter-regional synchrony throughout sleep (Figure 2d), consistent with reports that NREM sleep contains both highly coordinated and locally restricted slow waves (Bernardi et al., 2018).

### 2.3. Selective synaptic loss differentially modulates slow oscillation amplitude

To model age-related structural degradation, we systematically removed synapses from specific connection classes and quantified the resulting changes in SO peak-to-peak amplitude (Figure 3).

**Figure 3.**
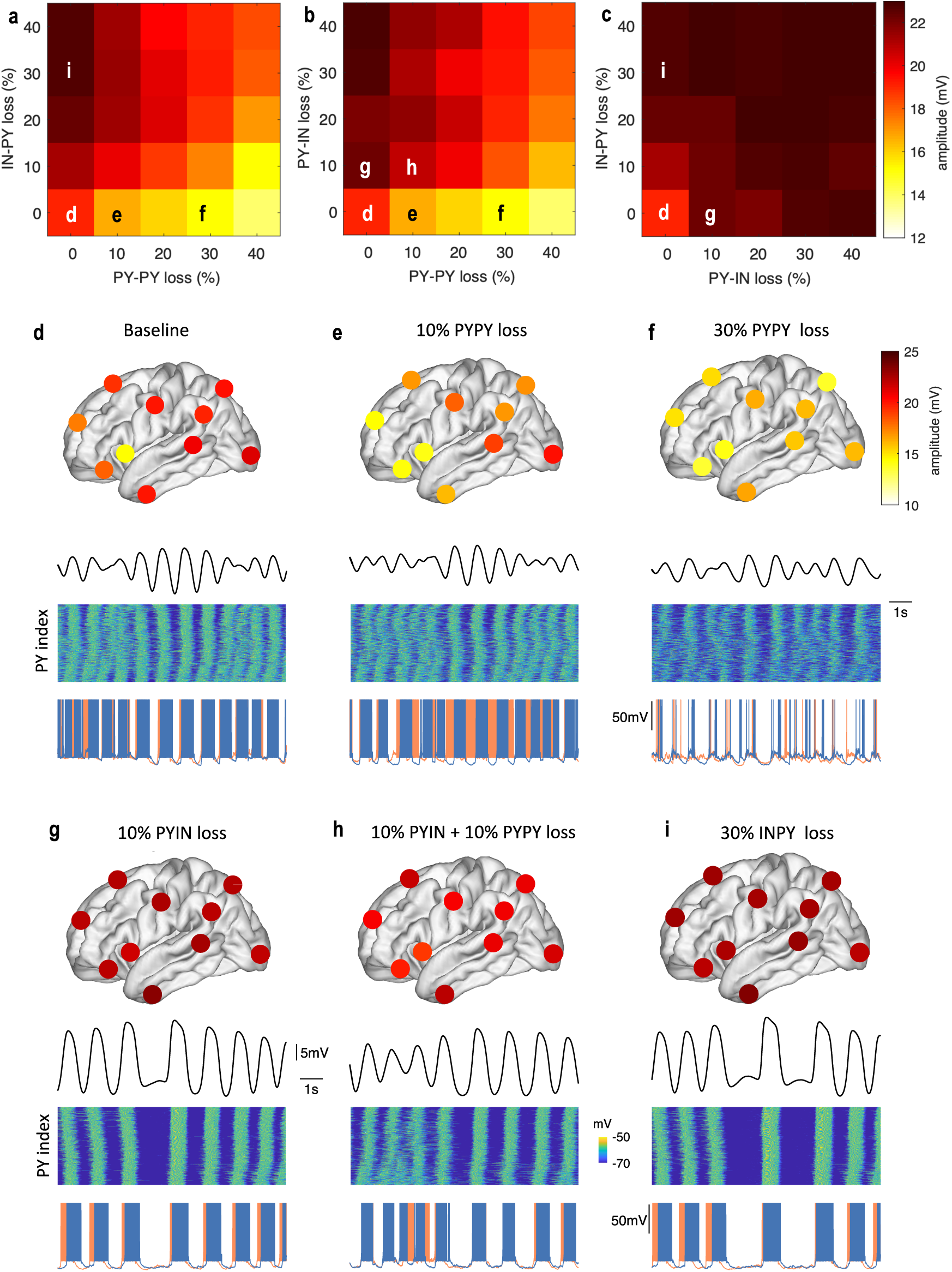
a) Average SO amplitude for different combinations of percentage of IN-PY vs PY-PY connections lost. b) As a), for PY-IN vs PY-PY. c) As (a), for IN-PY vs PY-IN. d) Average SO amplitude in 10 local (5mm radius) cortical regions for the baseline model. From top to bottom: regional SO amplitudes on the cortical surface, average L2 voltage, heatmap of L2 voltages, two pyramidal cell traces. e-f) As (d), for 10% and 30% PY-PY connections lost. SOs become less regular with smaller amplitude with PY-PY loss. g-i) As (d), for 10% PY-IN loss, 10% PY-IN + 10% PY-PY loss, and 30% IN-PY loss. Up-states are mode regular with higher amplitude.

Selective removal of recurrent excitatory PY-PY synapses produced a monotonic reduction in SO amplitude across the cortex (Figure 3d-f). In contrast, removal of inhibitory-to-excitatory (IN-PY) synapses increased SO amplitude. Manipulations of PY-IN projections yielded non-monotonic effects; moderate PY-IN loss increased SO amplitude, while extreme removal reduced it. Similarly, joint removal of PY–PY and PY–IN connections increased amplitude under moderate levels of degradation, inconsistent with the age-related reduction in SO amplitude observed experimentally.

Spatially, the baseline model exhibited regional heterogeneity, with larger SO amplitudes in posterior cortical regions (Figure 3d). Selective PY–PY loss reduced amplitude throughout the cortex (Figure 3f), whereas combined PY–PY and PY–IN removal increased amplitude and produced more regular oscillatory activity (Figure 3h). Because healthy aging is unlikely to involve synaptic loss substantially exceeding 30%, we focused subsequent analyses on a 30% reduction in PY–PY connectivity as a biologically plausible aged condition.

### 2.4. PY-PY loss reproduces age-related EEG changes in SO morphology

To determine whether selective PY–PY degradation reproduces age-related changes beyond SO amplitude, we quantified additional SO metrics previously used in human sleep studies (Djonlagic et al., 2021), and compared them with EEG recordings from different age cohorts (Figure 4).

**Figure 4.**
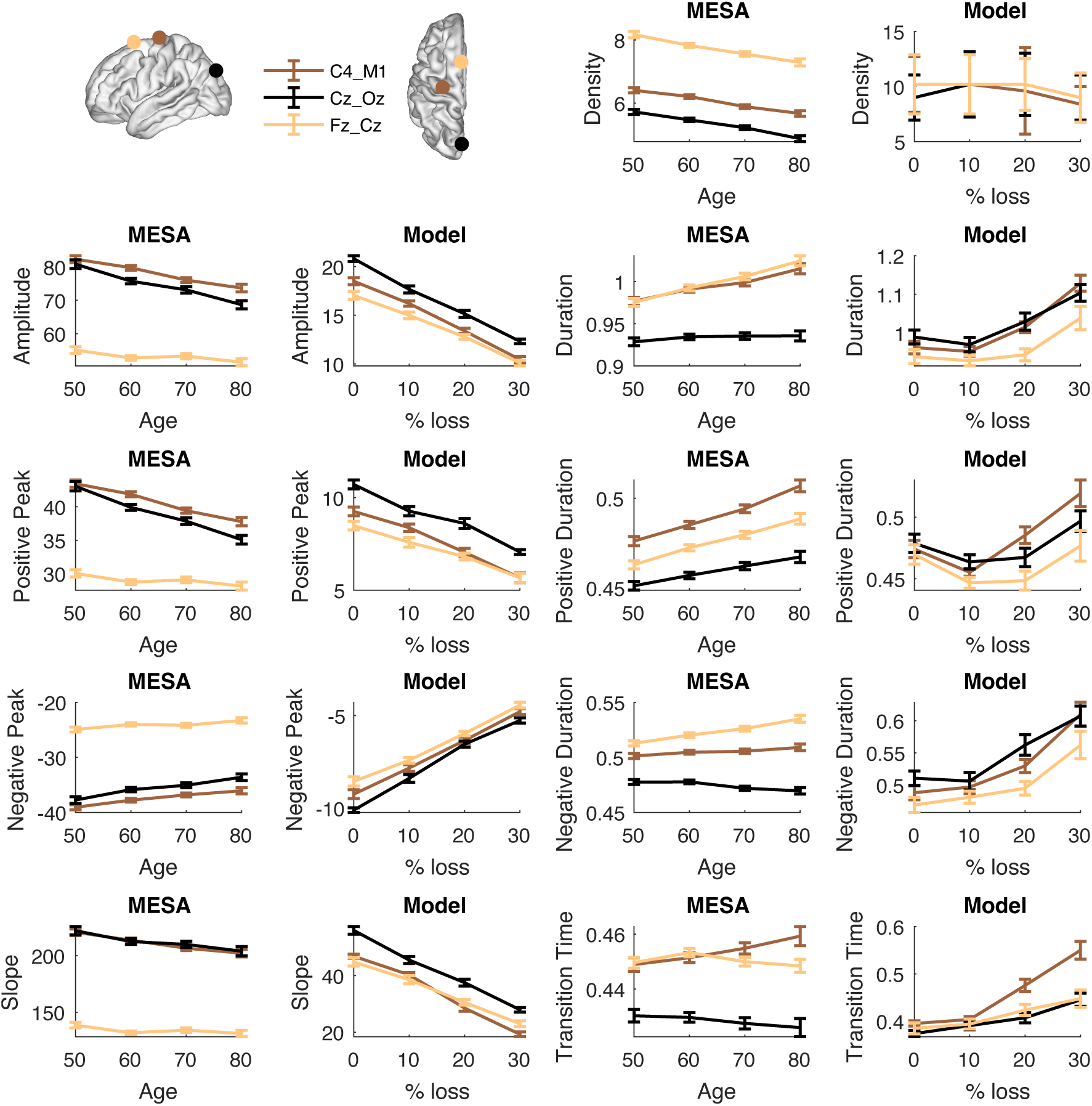
SO properties for three empirical and simulated EEG channels. a) 3D representation of channel locations. Two datasets are combined: one with recordings from a single EEG channel C4_M1, and one with two derivations Oz_Cz and Cz-Fz. Model channels are simulated by averaging the activities of neurons in a broad area around the approximate locations of C3 (the left-hemisphere analog of C4), Fz, and Oz. b) SO density (count per minute) for each channel in the empirical MESA recordings for ages 50 through 80 (left) and the model with for 0-30% PY-PY connections removed (right). c-i) As b, for MESA and Model SO amplitude, duration, positive peak amplitude, positive-half-wave duration, negative peak amplitude, negative-half-wave duration, slope, and transition time (time between negative and positive peaks). Similar trends with aging and synaptic loss are observed in model and empirical SOs. Data from (Djonlagic et al., 2021).

The empirical dataset combined measurements from MESA C4 recordings with frontal (Fz–Cz) and central-occipital (Cz–Oz) derivations from an independent cohort. To enable comparison across spatial scales, we generated three model EEG channels by averaging neuronal activity within broad cortical areas corresponding approximately to the empirical electrode locations.

Progressive PY–PY loss produced monotonic decreases in SO amplitude, density, and down-to-up transition slope, accompanied by increased SO duration. The model further reproduced reductions in both positive and negative peak amplitudes and increases in positive- and negative-half-wave durations (particularly for central and frontal electrodes in the latter) observed in the human EEG data.

A notable correspondence emerged in SO transition time, defined as the interval between negative and positive SO peaks. In the empirical data, transition times increased with age in central electrodes, whereas frontal electrodes followed a more complex trajectory, initially overlapping with central measurements before diverging at older ages. The model reproduced this pattern: transition times were similar across channels at low levels of synaptic loss but diverged progressively with increasing PY–PY degradation, with the central channel exhibiting the strongest increase.

Overall, the model captured the principal age-related trends reported in EEG studies, including reduced SO density and amplitude together with longer SO waveforms (Carrier et al., 2011; Djonlagic et al., 2021; Mander et al., 2013).

### 2.5. Spatial organization of EEG SO properties with aging

Beyond reproducing age-related trends in individual SO metrics, the model also captured several aspects of their spatial organization. This was particularly evident for amplitude-related measures, where the model reproduced the larger total, positive, and negative peak amplitudes observed in central and occipital channels relative to frontal channels. Similarly, steeper SO slopes and shorter transition times in posterior channels were consistent with empirical observations.

Duration-related metrics showed stronger agreement in their age-dependent trends than in their spatial organization. Nevertheless, the model reproduced the increased positive-half-wave duration observed in central electrodes with respect to other locations at advanced levels of synaptic loss.

For several measures, including SO density and duration, agreement with empirical data improved after approximately 10% PY–PY loss. Because the youngest empirical cohort was approximately 50 years old and measurable synaptic loss is expected to begin after the age of 40 (Markov et al., 2022), we designated the 10% PY–PY condition as corresponding to the youngest cohort. With this mapping, we directly compared young and aged model states at finer spatial resolution using activity sampled from ten local cortical regions with a radius of 5mm, comparable to LFP recordings.

Figure 5 illustrates both local activity traces and the spatial distribution of SO properties. In the young model, SOs exhibited clear anterior-posterior gradients, with higher density in frontal regions and larger amplitudes in posterior regions. PY–PY degradation attenuated these spatial differences. The aged model displayed reduced frontal dominance and diminished regional heterogeneity, consistent with reports of weakened frontal slow-wave activity in aging (Carrier et al., 2011; Mander et al., 2013).

**Figure 5.**
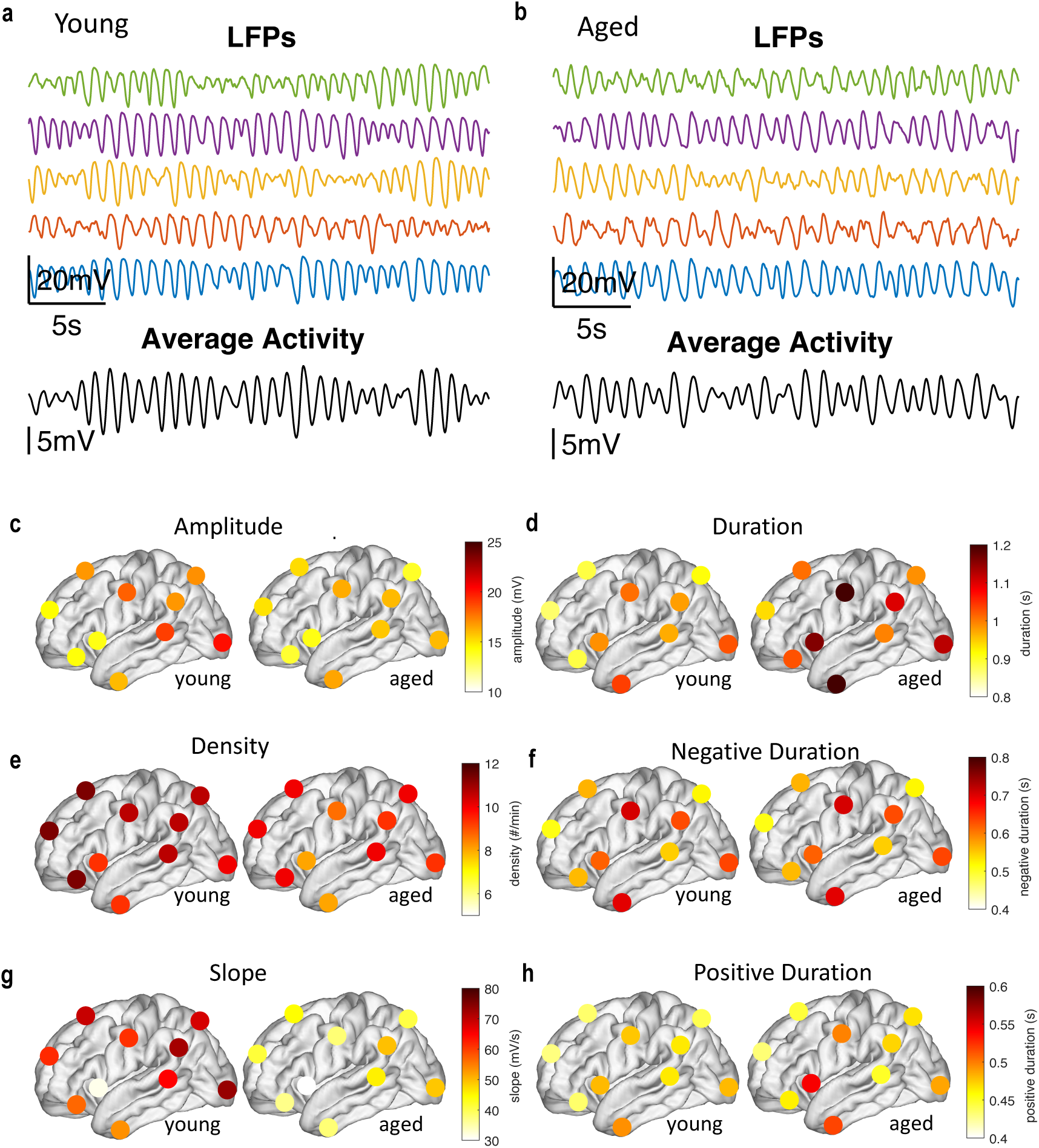
Young vs Aged model comparison. a) LFPs (top) and average volage activity (bottom) in the Young model. b) As a), for the Aged model (30% PY-PY synapses removed). c) Spatial distribution of amplitude from regions across the cortex in the young (left) and aged (right) models. d-h) As c), for SO duration, density, negative-half-wave duration, slope, and positive-half-wave duration.

At the EEG scale, SO duration increased with aging, primarily through elongation of the positive half-wave. While this trend agrees with both empirical data and previous literature, local LFP measurements showed a different pattern, with positive-half-wave duration remaining stable or decreasing. This discrepancy suggested that prolonged EEG-scale SOs might arise from changes in large-scale synchrony rather than longer local active states. To investigate this possibility, we examined neuronal dynamics at the single-cell level.

### 2.6. Age effects on network-level vs. single-cell level properties

Human EEG recordings showed an age-related increase in positive-half-wave duration, a trend reproduced by the model when activity was averaged across large cortical areas. In contrast, local LFP signals exhibited stable or decreasing positive-half-wave durations with increasing PY–PY loss. This divergence suggests that prolonged EEG-scale active states do not reflect longer local Up states, but instead emerge from reduced temporal coordination among cortical populations.

Analysis of single-cell activity supported this interpretation (Figure 6). Up states became shorter and more variable in the aged model (Figure 6a), and the number of spikes generated per Up state decreased substantially (Figure 6b). In addition, the phase-locking value between individual-cell Up states and the global cortical signal was reduced (Figure 6c), indicating weaker synchronization across the network.

**Figure 6.**
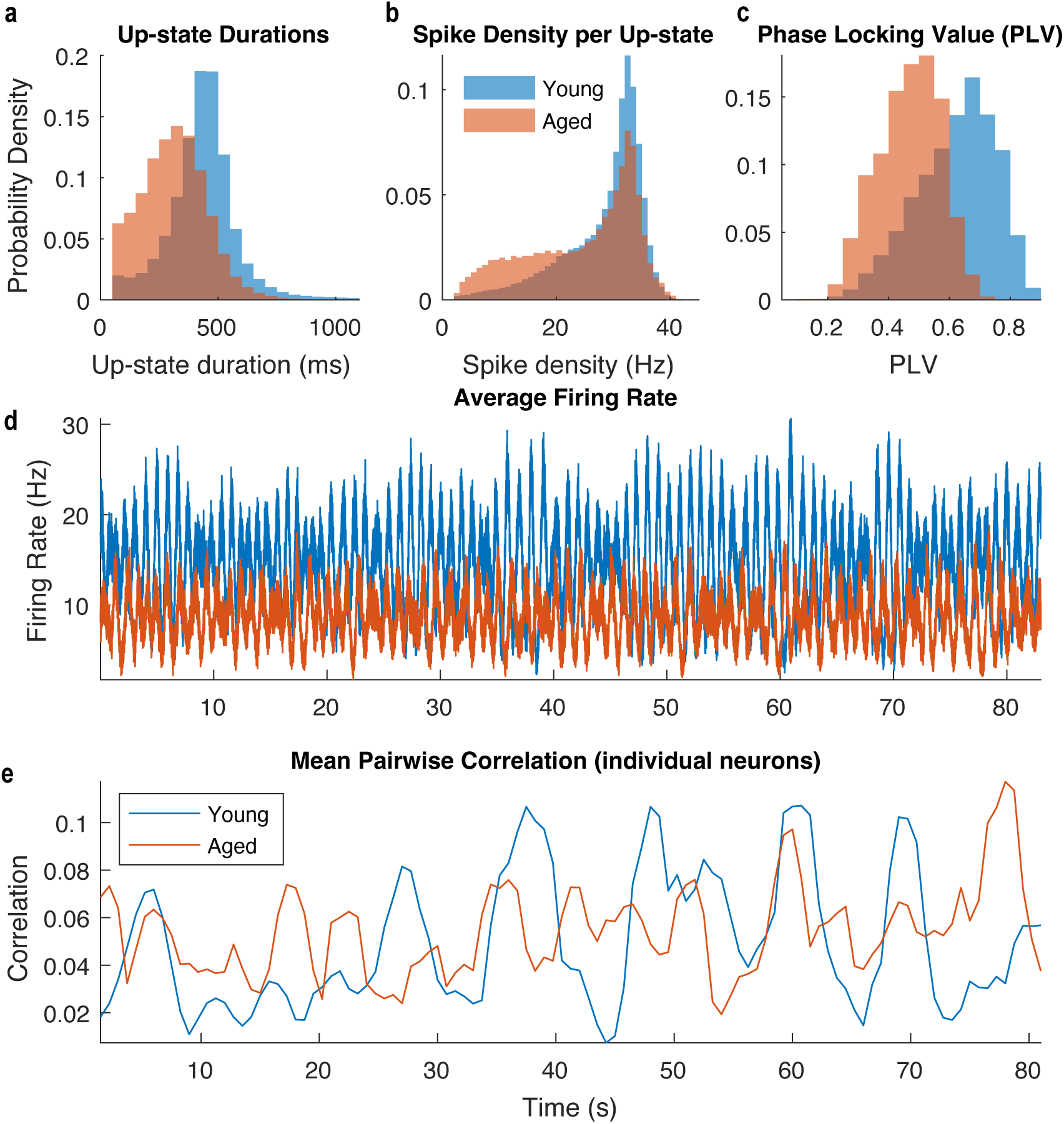
Individual cell properties in the Young vs Aged models. a) Distribution of up-state durations across individual pyramidal (PY) cells in cortical layer 2 (L2) for the Yound and Aged models. Only up-states with at least one spike are included. The distribution shifts to lower durations with age. b) Distribution of spike density (Hz) across all up-states for PY cells in L2. Spike density is defined as the rate of spiking per second in each up-state. The Aged model has more up-states with low spike densities. c) Phase locking value of each PY neuron in L2 with respect to the phase of the average voltage of all L2 PY cells. The distribution in the Aged model shifts towards lower phase locking values. d) Instantaneous firing rate over time for the Young and Aged models, showing lower overall firing rate for the Aged model. e) Mean pairwise correlation between 1000 random PY cells in L2 for the Young and Aged models. The Young model has more marked peaks of high correlation (indicative of synchrony) and clearer alternation between high and low synchrony periods.

Consistent with these observations, average cortical firing rates decreased with PY–PY degradation (Figure 6d). Pairwise correlations between neuronal membrane potentials also revealed fewer episodes of highly synchronized activity in the aged model. Periods of high correlation, indicative of high global synchrony, were more frequent and more pronounces in the young model and became less evident following synaptic loss.

Together, these results indicate that age-related prolongation of SOs at the EEG scale emerges from reduced synchrony among cortical populations rather than prolonged local Up states. At the cellular level, recurrent excitation loss produces shorter, weaker, and less coordinated Up states that become temporally dispersed across the cortex. When averaged over large spatial scales, these asynchronous events appear as broader, lower-amplitude active periods. This mechanism provides a potential link between structural brain aging, altered sleep oscillations, and impaired sleep-dependent plasticity and memory consolidation.

## 3. Discussion

In this study, we developed a multi-scale, whole-brain thalamocortical network model constrained by human structural connectivity to investigate how age-related synaptic degradation shapes slow-wave sleep (SWS) dynamics. The model reproduces key features of SWS in a baseline “young” condition and captures experimentally observed alterations in slow oscillation (SO) morphology under simulated aging. Specifically, selective degradation of recurrent excitatory pyramidal-to-pyramidal (PY–PY) connectivity reproduced the principal age-related changes observed in human EEG, including reduced SO amplitude, density, and slope, and increased oscillation duration.

Importantly, the model revealed that these macroscopic changes emerge despite shorter and less robust local Up states. Progressive PY–PY loss weakened local activity and reduced synchronization across cortical populations, causing temporally dispersed Up states that appear as broader, lower-amplitude oscillations at the EEG scale.

Our whole-hemisphere model integrates biologically grounded connectivity derived from diffusion MRI tractography of the Human Connectome Project and organized according to the HCP multimodal atlas. By mapping parcel-level connectivity statistics onto column-wise projections with distance-dependent delays, the model links large-scale structural constraints to emergent thalamocortical dynamics. This approach extends previous modeling studies that simulated aging as reduced cortical connectivity in smaller-scale networks (Wei et al., 2023), enabling investigation of how distributed synaptic degradation reshapes sleep activity across the cortex.

A central finding is that loss of recurrent excitation, rather than alterations in excitatory-inhibitory (PY-IN) pathways, best explains age-related changes in SO dynamics. Whereas removal of inhibitory-to-excitatory connections increased SO amplitude, selective PY–PY degradation consistently reduced SO amplitude and density across cortical locations. These results suggest that aging shifts cortical dynamics toward a state of reduced effective excitation. Rather than resulting primarily from increased inhibition, the observed SO changes are better explained by degradation of recurrent excitatory connectivity. Reduced recurrent excitation weakened local Up states, decreased spike output during active periods, and impaired synchronization among distant cortical populations. Together, these effects were sufficient to reproduce both local and large-scale signatures of aging observed in human sleep recordings.

This interpretation is consistent with anatomical evidence that aging is accompanied by reductions in dendritic arborization, spine density, and glutamatergic synaptic integrity (Burke & Barnes, 2006; Peters, 2002). It is also consistent with recent evidence linking structural brain integrity to sleep physiology. In a large study of approximately 600 participants from the Multi-Ethnic Study of Atherosclerosis (MESA), longer SO duration was associated with lower white-matter fractional anisotropy and reduced total white-matter volume (Kozhemiako et al., 2025). These findings were interpreted as reflecting impaired large-scale corticocortical connectivity rather than changes in local synaptic strength. Because long-range excitatory projections play a critical role in coordinating distributed cortical populations, their degradation would be expected to impair the propagation and synchronization of Up states, resulting in weaker and less coherent SOs.

The model also reproduced age-related changes in the spatial organization of slow oscillations. In the baseline condition, SO density was highest in frontal cortex, consistent with the well-established anterior predominance of slow-wave activity in young adults (Riedner et al., 2007) These spatial gradients became less pronounced following PY–PY degradation, leading to reduced regional heterogeneity. This finding aligns with reports of diminished frontal slow-wave activity and reduced topographic differentiation in older adults (Helfrich et al., 2018; Mander et al., 2013), and suggests that progressive degradation of recurrent cortical connectivity contributes to the loss of regional specialization observed with aging.

A notable finding was the dissociation between local and global measures of SO dynamics. At the EEG scale, both empirical data and model activity exhibited longer SO durations and prolonged positive half-waves with aging. In contrast, local field potentials and single-neuron activity revealed the opposite trend: Up states became shorter, weaker, and more variable as PY–PY connectivity was reduced. The model reconciles these observations by demonstrating that recurrent excitation loss reduces synchronization across cortical populations. As local Up states become less phase-locked, their temporal overlap decreases, generating broader active periods when activity is averaged across large spatial scales. Thus, age-related prolongation of SOs in EEG recordings may not reflect longer cortical Up states, but rather a loss of temporal coordination among neuronal populations.

This interpretation is supported by experimental evidence that aging is accompanied by reduced large-scale synchronization of sleep oscillations and weaker coupling between cortical slow oscillations and other sleep rhythms (Helfrich et al., 2018; Muehlroth et al., 2019). Our results provide a potential circuit-level mechanism linking these observations to structural degradation of recurrent cortical excitation.

These findings have important implications for understanding cognitive aging. Slow oscillations provide the temporal framework that coordinates thalamic spindles and hippocampal sharp-wave ripples during memory consolidation (Diekelmann & Born, 2010; Klinzing et al., 2019). Reductions in SO amplitude and slope are associated with impaired memory retention in older adults (Mander et al., 2013). The present results suggest that aging may impair this coordinating function through two complementary mechanisms. First, recurrent excitation loss reduces the strength of local Up states and the number of spikes generated during active periods, potentially limiting opportunities for synaptic plasticity. Second, reduced synchronization across cortical populations broadens and disperses active states in time, potentially weakening the precise temporal relationships required for cross-regional communication during sleep. Together, these mechanisms provide a plausible link between structural brain aging, altered sleep oscillations, and age-related memory decline.

Recent evidence from a parallel rodent study further supports this interpretation. In that work, shorter Up states were sufficient to consolidate newly acquired hippocampus-dependent memories but failed to preserve older cortical memories (Golden et al., 2025). Together with the present findings, these results suggest that age-related restructuring of SO dynamics may disrupt interference-free memory consolidation during NREM sleep and contribute to the memory deficits observed in older adults.

Several limitations should be noted. Aging was modeled as uniform synaptic weakening or removal within specific connection classes, whereas biological aging likely involves heterogeneous, cell-type-specific, and regionally variable changes in synaptic density, neuromodulation, intrinsic excitability, and white-matter integrity. Future work could incorporate age-specific tractography datasets, layer-specific degeneration patterns, and activity-dependent plasticity mechanisms to better capture the progression of age-related cognitive decline.

In summary, selective loss of recurrent cortical excitation was sufficient to reproduce age-related changes in slow-oscillation amplitude, density, slope, duration, and spatial organization. Beyond reproducing these empirical observations, the model provides a mechanistic explanation for their emergence across scales. Recurrent excitation loss weakens local Up states and reduces synchronization among cortical populations, transforming short, coordinated activity bursts into temporally dispersed network events. These results establish a direct link between structural brain aging and altered sleep dynamics, and identify recurrent cortical excitation as a potential target for interventions aimed at preserving sleep-dependent cognitive function throughout aging.

## 4. Methods

### 4.1. Model neurons

#### 4.1.1. Map-based cortical neurons

The model has 10,242 cortical columns uniformly positioned across the surface of one hemisphere, one for each vertex in the ico5 cortical reconstruction previously reported in Rosen et al. (2019). The medial wall includes 870 of these vertices, so all analyses for this one-hemisphere model were done on the remaining 9372 columns. Each column has 6 cortical layers (L2, L3, L4, L5a, L5b and L6). For each layer in each column, one excitatory (PY) and one inhibitory (IN) neuron were simulated. To allow for large-scale network simulations we modeled cortical neurons using a “map” model based on difference equations (Rulkov, 2002; Rulkov et al., 2004).

Map-based models sample membrane potential using a large discrete time step compared to the widely used conductance-based models described by ordinary differential equations and still capture neural responses from these models. Importantly, map-based models replicate spiking activity of cortical fast-spiking, regular-spiking and intrinsically bursting neurons (Rulkov et al., 2004). Variations of the map model have enabled the simulation of large-scale brain networks with emergent oscillatory activity (Rulkov & Bazhenov, 2008), including realistic sleep spindles (Rosen et al., 2019) and cortical slow oscillations (Komarov et al., 2018). In our study, we used the previously proposed map model modification (Komarov et al., 2018), which implemented nonlinear dynamical bias, activity dependent depolarizing mechanisms, and slow hyperpolarizing mechanisms capturing the effects of leak currents, the Ca2+ dependent K+ currents and the persistent Na+ currents, which were found to be critical for simulating up/down state transitions during SO. Parameter values were initially set to the previously reported reference values (Komarov et al., 2018), which were shown to produce biologically realistic neural behavior during SO, and then tuned with small variations to maintain adequate neural responses in our comparatively larger-scale network. The map model equations are described below, and all parameter values are reported in (Marsh et al., 2024).

The activity of individual pyramidal PY cells was described in terms of four continuous variables sampled at discrete moments of time *n* : *x*_*n*_, dictating the trans-membrane potential *V*_*n*_ = 50*x*_*n*_ − 15; *y*_*n*_, representing slow ion channel dynamics; *u*_*n*_, describing slow hyperpolarizing currents; and *k*_*n*_, determining the neuron sensitivity to inputs. The activity of IN cells, which biologically have faster spiking dynamics, was modeled using only variable *x*_*n*_ and fixing the other variables to constant values. Dynamical PY variables evolve according to the following system of difference equations, where *n* = 1 , 2, … indexes time steps of 0.5ms in size (Rulkov & Bazhenov, 2008).

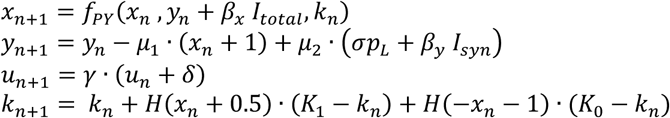

where function *H*(*w*) denotes the Heaviside step function, which takes the value 0 for *w* ≤ 0 and 1 otherwise. The reduced model for IN cells is given by

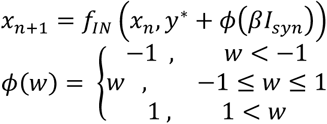

where *y*^∗^ is a fixed value of *y*_*n*_. We describe each of the system’s components below.

**Spike-generating Systems**: Variables *x*_*n*_ and *y*_*n*_ model the fast neuron dynamics, representing the effects of the fast Na+ and K+ currents responsible for spike generation. The nonlinear function *f*_*PY*_, modified by (Komarov et al., 2018) from (Rulkov, 2002), determines the spike waveform as

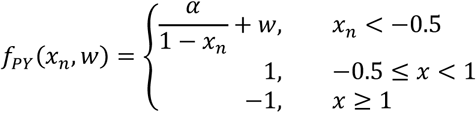

where *α* is a constant parameter that controls the non-linearity of the spike-generating mechanism. Parameters *μ*_1_, *μ*_2_ < 1 modulate the slower updating of variable *y* with respect to variable *x*, and parameter *σ* dictates the neuron resting potential as (1-*σ*). The scaling constants *β*_*x*_, *β*_*y*_ modulate the input current *I*_*syn*_ for variables *x* and *y*, respectively. The neuron’s membrane potential is calculated as *V*_*n*_ = 50*x*_*n*_ − 15 and a spike is generated if and only if *V*_*n*_ > 0.1. In simulations, a lower bound of 10^−4^ was manually imposed for the quantity *β*_*x*_*I*_*total*_.

For inhibitory neurons IN, function *f*_*IN*_ is given by

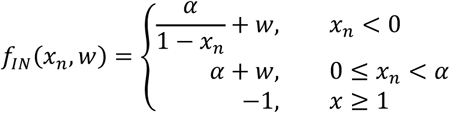

A spike is generated if and only if 0 < *x*_*n*_ ≤ *α* + *y*^∗^ + *ϕ*(*β*_*x*_*I*_*total*_) and *x*_*n*–1_ ≤ 0.

**Adaptation Equation:** The slow variable *u*_*n*_ represents the effects of the slow hyperpolarizing potassium currents (*I*_*D*_, explained below) reducing neural excitability over the course of the Up state and involved in Up state termination. The value of *γ* is taken as 0.99 for *x*_*n*_ < −1 and 0.995 otherwise. Parameter *δ*= 0.24 has an effect only if *x*_*n*_ < *x*_*n*_ < 1 and is set to 0 othewise.

**Sensitivity Equation:** The variable *k*_*n*_ reduces neuron sensitivity to inputs during Up states (i.e. during high spiking activity) by regulating the spike nonlinearity function *f*_*P*F_. This gives a net phenomenological representation of the combined effects of high-level synaptic activity, increase in voltage-gated hyperpolarizing currents, and/or adaptation of fast Na+ channels.

**Computation of currents**: The total current *I*_*total*_ is a sum of external applied DC current *I*_*DC*_, synaptic inputs ( *I*_*syn*_), persistent Na+ current ( *I*_*syn*_) slow hyperpolarizing currents ( *I*_*D*_), and leak currents ( *I*_*leak*_):

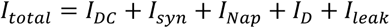

For each neuron *i*, the synaptic current *I*_*syn*_ is the sum of individual AMPA, GABA_A_ and NMDA synaptic currents. The individual synaptic current from neuron *j* to neuron *i* has the general form

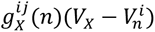

for AMPA and GABA synapses, and

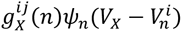

for NMDA synapses, with an additional modulator *ψ*_*n*_. Here *g*_*x*_^*ij*^ (*n*) represents conductance at time n. See (Marsh et al., 2024) for details.

Lastly, miniature post-synaptic potentials are only present in AMPA and GABA synapses. They are modeled as spontaneous events from a random Poisson process that increase conductance, which produces miniature post-synaptic potentials (“minis”) (Stevens, 1993). The frequency is modulated in time to account for mini reduction after Up-states. See (Marsh et al., 2024) for details.

#### 4.1.2. Hodgkin-Huxley thalamic neurons

The thalamus was modeled using a network of core (specific) and matrix (non-specific) nuclei, each consisting of thalamic relay (TC) and reticular (RE) neurons. We simulated 642 thalamic cells of each of these four types. TC and RE cells were modeled as single compartments with membrane potential V governed by the Hodgkin-Huxley kinetics, so that the total membrane current per unit area is given by

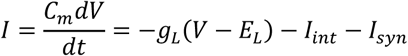

where *C*_*m*_is the membrane capacitance, *g*_*L*_ is the non-specific (mixed Na+ and Cl-) leakage conductance, *E*_*L*_ is the reversal potential, *I*_*int*_ is a sum of active intrinsic currents, and *I*_*syn*_ is a sum of synaptic currents.

Thalamic reticular and relay neurons included intrinsic properties necessary for generating rebound responses which were found to be critical for spindle generation. Intrinsic currents for both RE and TC cells included a fast sodium current, a fast potassium current, a low-threshold Ca2+ “T” current, and a potassium leak current. For TC cells, an additional hyperpolarization-activated cation “h” current was also included. See (Marsh et al., 2024) for details on intrinsic and synaptic thalamic currents.

#### 4.1.3. Stage Transitions

The transition from wake to slow wave sleep involved increasing the strength of the slow hyperpolarizing currents *I*_*D*_ (similar to a calcium current) and the potassium leak currents *I*_*leak*_. We further increased the strength of AMPA and GABA synaptic currents, as well as the strength of TC potassium leak currents, based on (Krishnan et al., 2016).

### 4.2. Network Structure

Our whole hemisphere computational model has realistic local and long-range cortical connectivity. Connections within the column follow those described in the canonical cortical circuit (Fig 1a). Additionally, connections within 0.1 mm are strengthened by a factor of 5 to mimic in-vivo increased strength of proximal synapses (Hawkins & Ahmad, 2016; London & Segev, 2001). Inhibition is local: IN cells project only to PY cells in the same layer, with connections only in their own column, 1st and 2nd degree neighbors. The strength of inhibitory synapses is 2 times larger within the local column than outside. Finally, miniature postsynaptic potentials on intracortical AMPA synapses are modeled as a Poisson process, as in (Krishnan et al., 2018).

Thalamocortical (TC) and reticular (RE) neurons are modeled using Hodgkin-Huxley dynamics (Hodgkin & Huxley, 1952), with 2 subtypes of each to represent matrix and core neurons. The matrix TC neurons have a 45 mm (and 80 mm) fanout radius to cortical excitatory (and inhibitory) neurons while core TC neurons have 12 mm and 13 mm radii, respectively. Within-thalamic connections have 11 mm radii. There are 642 neurons of each of these 4 types (thalamic matrix, thalamic core, reticular matrix, reticular core).

#### 4.2.1. Hierarchically Guided Laminar Connectivity

Long-range connectivity in the model is primarily based on the cortical parcellation into 180 areas proposed by the Human Connectome Project (HCP)-MMP1.0 atlas (Glasser et al., 2016; Van Essen et al., 2013). Each parcel was assigned a hierarchical index inversely proportional to its estimated bulk myelination (Burt et al., 2018; Glasser & Essen, 2011). Excitatory cortical connections (PY-PY) were then split into 6 classes according to the pre- and post-synaptic hierarchical index: within the local column, within the local parcel (or between contralateral homologs), strongly (>50th percentile) or weakly ≤50th percentile) feedforward (from lower to higher hierarchical index), and strongly or weakly (>, ≤ 50th percentile respectively) feedback (from higher to lower hierarchical index). Based on previous connectivity reports (Markov et al., 2013; Rockland, 2019), synaptic weights for these connections were scaled by the factors shown in Fig 1c. Note that, while these factors impose constraints on the relative inter-layer connection strengths, the absolute value of synaptic strength is a free parameter.

#### 4.2.2. Diffusion MRI Guided Connection Probability

The connection probability was further scaled based on previous diffusion MRI (dMRI) connectivity studies (Rosen & Halgren, 2021). These observations reveal an exponential decay relationship between inter-parcel dMRI connectivity and fiber distance. To obtain a probability of connection at the scale of columns, approximate intercolumnar fiber distances were estimated from geodesic distances. For parcel centroids, intra-hemispheric dMRI streamline distances and their corresponding geodesic distances were found to be related to a first approximation by a linear rational function. Column-wise fiber distances were estimated by applying this function to intercolumnar geodesic distances. Thus, a distance-dependent probability of connection between columns was obtained by applying the dMRI exponential relation to column-wise fiber distances. Moreover, the residual parcel-wise dMRI connectivity (not distance related, obtained by regressing out the exponential trend) was added back to intercolumnar connections based on the columns’ parcels. This distance-independent connectivity accounts for the functional specialization of each parcel. Lastly, conduction delays were set to be proportional to fiber tract distances, with the longest connection (226mm) having an assigned delay of 40ms.

### 4.3. Simulating Effects of Aging

Age was simulated as synaptic loss from the baseline model connectivity. To determine the specific type of synaptic loss that best captured age effects, we first conducted sweeps with all combinations of 0-40% synaptic loss of PY-PY, PY-IN, and IN-PY connections. Synaptic loss of X% was implemented as setting the strength of each connection to 0 with X% probability (e.g. each synapse was removed with 30% probability in the 30% loss simulation).

### 4.4. Sleep Properties

#### 4.4.1. Channel signals

A channel signal in the model was obtained by averaging the membrane voltages of all PY and IN cells from all six layers within a radius of 5mm for LFP signals and 150mm for EEG signals. The 10 LFP channel locations were chosen so that each region would belong to one of the 10 functional networks identified in (Rosen & Halgren, 2021), to favor independence between channels.

The 3 EEG channel locations in the model were selected at the approximate locations of EEG electrodes C3, Fz, and Oz. Because the model includes only the left hemisphere, the central electrode was situated as C3, although the empirical data comes from the analogous right-hemisphere recording (C4).

#### 4.4.2. Slow-Oscillation Detection

SO detection in the model was adapted from the methodology in (Djonlagic et al., 2021). Each channel was detrended and filtered in the 0.3-4Hz band. The positive-to-negative zero crossings were then marked. Intervals between zero-crossings of at least 0.5 s in duration were marked. SOs were then detected using a relative minimal amplitude threshold. The threshold for peak-to-peak amplitude was the 85^th^ percentile of peak-to-peak amplitudes in the channel, and the threshold for negative (down-state) amplitude was the 40^th^ percentile of negative-peak amplitudes in the channel.

#### 4.4.3. Slow-Oscillation Properties

SO amplitude was computed as the peak-to-peak channel amplitude. Positive amplitude is the value of the positive peak, and negative amplitude is the value of the negative peak. Density was estimated by binning SO events into 10s time intervals covering the whole simulation time and averaging across time intervals. Duration was the time difference between the SO zero-crossings. The total duration was split into negative and positive halfwave durations at the negative-to-positive zero-crossing. The negative-to-positive slope was obtained as the SO peak-to-peak amplitude divided by the time between negative and positive peaks. The transition time was the time between negative and positive peaks.

#### 4.4.4. Individual Cell Up-states and Properties

For individual cell property analysis, only cells in L2 and not belonging to the medial wall were considered. For Up-state detection, voltages were thresholded above −60mV to obtain a binary up-state vs. not-up-state vector. Up-state periods closer than 50ms were merged and up-state periods shorter than 50ms were removed. Up-states without any spikes were excluded.

Spike density per up-state was obtained by counting the number of spikes in each up-state and dividing by the up-state duration in seconds. Phase locking values for each cell were computed for each cell with respect to the average voltage signal of all L2 cells. Mean pairwise correlation over time was calculated for 3s windows across 1000 randomly sampled L2 PY cells.

### 4.5. Empirical Data

Empirical SO properties were obtained from EEG recordings in the MESA age dataset in (Djonlagic et al., 2021). All properties are reported as mean +- SEM (standard error of the mean).

## Notes

### Competing Interest Statement

The authors have declared no competing interest.

### Summary of Updates

This version of the manuscript was revised to update the results. Specifically, new figures 4 and 6 were included.

### Funding

- NIH: 1R01AG099626, 1RF1NS132913, 1R01MH125557

## References

Bazhenov, M., Timofeev, I., Steriade, M., & Sejnowski, T. J. (1998a). Cellular and Network Models for Intrathalamic Augmenting Responses During 10-Hz Stimulation. Journal of Neurophysiology, 79(5), 2730–2748. 10.1152/jn.1998.79.5.2730

Bazhenov, M., Timofeev, I., Steriade, M., & Sejnowski, T. J. (1998b). Computational Models of Thalamocortical Augmenting Responses. Journal of Neuroscience, 18(16), 6444–6465. 10.1523/JNEUROSCI.18-16-06444.1998

Bernardi, G., Siclari, F., Handjaras, G., Riedner, B. A., & Tononi, G. (2018). Local and Widespread Slow Waves in Stable NREM Sleep: Evidence for Distinct Regulation Mechanisms. Frontiers in Human Neuroscience, 12. https://www.frontiersin.org/articles/10.3389/fnhum.2018.00248

Bonjean, M., Baker, T., Lemieux, M., Timofeev, I., Sejnowski, T., & Bazhenov, M. (2011). Corticothalamic Feedback Controls Sleep Spindle Duration In Vivo. Journal of Neuroscience, 31(25), 9124–9134. 10.1523/JNEUROSCI.0077-11.2011

Burke, S. N., & Barnes, C. A. (2006). Neural plasticity in the ageing brain. Nature Reviews Neuroscience, 7, 30–40.

Burt, J. B., Demirtaş, M., Eckner, W. J., Navejar, N. M., Ji, J. L., Martin, W. J., Bernacchia, A., Anticevic, A., & Murray, J. D. (2018). Hierarchy of transcriptomic specialization across human cortex captured by structural neuroimaging topography. Nature Neuroscience, 21(9), 1251–1259. 10.1038/s41593-018-0195-0

Carrier, J., Viens, I., Poirier, G., Robillard, R., Lafortune, M., Vandewalle, G., Martin, N., Barakat, M., Paquet, J., & Filipini, D. (2011). Sleep slow wave changes during the middle years of life. European Journal of Neuroscience, 33(4), 758–766. 10.1111/j.1460-9568.2010.07543.x

Diekelmann, S., & Born, J. (2010). The memory function of sleep. Nature Reviews Neuroscience, 11(2), Article 2. 10.1038/nrn2762

Djonlagic, I., Mariani, S., Fitzpatrick, A. L., Van Der Klei, V. M. G. T. H., Johnson, D. A., Wood, A. C., Seeman, T., Nguyen, H. T., Prerau, M. J., Luchsinger, J. A., Dzierzewski, J. M., Rapp, S. R., Tranah, G. J., Yaffe, K., Burdick, K. E., Stone, K. L., Redline, S., & Purcell, S. M. (2021). Macro and micro sleep architecture and cognitive performance in older adults. Nature Human Behaviour, 5(1), 123–145. 10.1038/s41562-020-00964-y

Dubé, J., Lafortune, M., Bedetti, C., Bouchard, M., Gagnon, J. F., Doyon, J., Evans, A. C., Lina, J.-M., & Carrier, J. (2015). Cortical Thinning Explains Changes in Sleep Slow Waves during Adulthood. Journal of Neuroscience, 35(20), 7795–7807. 10.1523/JNEUROSCI.3956-14.2015

Glasser, M. F., Coalson, T. S., Robinson, E. C., Hacker, C. D., Harwell, J., Yacoub, E., Ugurbil, K., Andersson, J., Beckmann, C. F., Jenkinson, M., Smith, S. M., & Van Essen, D. C. (2016). A multi-modal parcellation of human cerebral cortex. Nature, 536(7615), 171–178. 10.1038/nature18933

Glasser, M. F., & Essen, D. C. V. (2011). Mapping Human Cortical Areas In Vivo Based on Myelin Content as Revealed by T1- and T2-Weighted MRI. Journal of Neuroscience, 31(32), 11597–11616. 10.1523/JNEUROSCI.2180-11.2011

Hawkins, J., & Ahmad, S. (2016). Why Neurons Have Thousands of Synapses, a Theory of Sequence Memory in Neocortex. Frontiers in Neural Circuits, 10, 23. 10.3389/fncir.2016.00023

Helfrich, R. F., Mander, B. A., Jagust, W. J., Knight, R. T., & Walker, M. P. (2018). Old Brains Come Uncoupled in Sleep: Slow Wave-Spindle Synchrony, Brain Atrophy, and Forgetting. *Neuron*, 97, 221–230 224,. 10.1016/j.neuron.2017.11.020

Hodgkin, A. L., & Huxley, A. F. (1952). A quantitative description of membrane current and its application to conduction and excitation in nerve. The Journal of Physiology, 117(4), 500–544. 10.1113/jphysiol.1952.sp004764

Klinzing, J. G., Niethard, N., & Born, J. (2019). Mechanisms of systems memory consolidation during sleep. Nature Neuroscience, 22, 1598–1610,. 10.1038/s41593-019-0467-3

Komarov, M., Krishnan, G., Chauvette, S., Rulkov, N., Timofeev, I., & Bazhenov, M. (2018). New class of reduced computationally efficient neuronal models for large-scale simulations of brain dynamics. Journal of Computational Neuroscience, 44(1), 1–24. 10.1007/s10827-017-0663-7

Kozhemiako, N., Heckbert, S. R., Castro-Diehl, C., Paquet, C. B., Bertisch, S. M., Habes, M., Fohner, A. E., Bryan, R. N., Nasrallah, I., Hughes, T. M., Redline, S., & Purcell, S. M. (2025). Mapping the relationships between structural brain MRI characteristics and sleep electroencephalography patterns: The Multi-Ethnic Study of atherosclerosis. Sleep, 48(8), zsaf074. 10.1093/sleep/zsaf074

Krishnan, G. P., Chauvette, S., Shamie, I., Soltani, S., Timofeev, I., Cash, S. S., Halgren, E., & Bazhenov, M. (2016). Cellular and neurochemical basis of sleep stages in the thalamocortical network. eLife, 5, e18607. 10.7554/eLife.18607

Krishnan, G. P., Rosen, B. Q., Chen, J.-Y., Muller, L., Sejnowski, T. J., Cash, S. S., Halgren, E., & Bazhenov, M. (2018). Thalamocortical and intracortical laminar connectivity determines sleep spindle properties. PLOS Computational Biology, 14(6), e1006171. 10.1371/journal.pcbi.1006171

London, M., & Segev, I. (2001). Synaptic scaling in vitro and in vivo. Nature Neuroscience, 4(9), Article 9. 10.1038/nn0901-853

Mander, B. A., Rao, V., Lu, B., Saletin, J. M., Lindquist, J. R., Ancoli-Israel, S., Jagust, W., & Walker, M. P. (2013). Prefrontal atrophy, disrupted NREM slow waves and impaired hippocampal-dependent memory in aging. Nature Neuroscience, 16(3), 357–364. 10.1038/nn.3324

Markov, N. T., Ercsey-Ravasz, M., Van Essen, D. C., Knoblauch, K., Toroczkai, Z., & Kennedy, H. (2013). Cortical High-Density Counterstream Architectures. Science, 342(6158), 1238406. 10.1126/science.1238406

Markov, N. T., Lindbergh, C. A., Staffaroni, A. M., Perez, K., Stevens, M., Nguyen, K., Murad, N. F., Fonseca, C., Campisi, J., Kramer, J., & Furman, D. (2022). Age-related brain atrophy is not a homogenous process: Different functional brain networks associate differentially with aging and blood factors. Proceedings of the National Academy of Sciences, 119(49), e2207181119. 10.1073/pnas.2207181119

Marsh, B., Navas-Zuloaga, M. G., Rosen, B. Q., Sokolov, Y., Delanois, J. E., Gonzalez, O. C., Krishnan, G. P., Halgren, E., & Bazhenov, M. (2024). Emergent effects of synaptic connectivity on the dynamics of global and local slow waves in a large-scale thalamocortical network model of the human brain. PLOS Computational Biology, 20(7), e1012245. 10.1371/journal.pcbi.1012245

Marshall, L., Helgadottir, H., Molle, M., & Born, J. (2006). Boosting slow oscillations during sleep potentiates memory. Nature, 444, 610–613.

Peters, A. (2002). Structural changes that occur during normal aging of primate cerebral hemispheres. Neuroscience & Biobehavioral Reviews, 26(7), 733–741. 10.1016/S0149-7634(02)00060-X

Riedner, B. A., Vyazovskiy, V. V., Huber, R., Massimini, M., Esser, S., Murphy, M., & Tononi, G. (2007). Sleep Homeostasis and Cortical Synchronization: III. A High-Density EEG Study of Sleep Slow Waves in Humans. Sleep, 30(12), 1643–1657.

Rockland, K. S. (2019). What do we know about laminar connectivity? NeuroImage, 197, 772–784. 10.1016/j.neuroimage.2017.07.032

Rosen, B. Q., C Halgren, E. (2021). A Whole-Cortex Probabilistic Diffusion Tractography Connectome. Eneuro, 8(1), ENEURO.0416-20.2020. 10.1523/ENEURO.0416-20.2020

Rosen, B. Q., Krishnan, G. P., Sanda, P., Komarov, M., Sejnowski, T., Rulkov, N., Ulbert, I., Eross, L., Madsen, J., Devinsky, O., Doyle, W., Fabo, D., Cash, S., Bazhenov, M., & Halgren, E. (2019). Simulating human sleep spindle MEG and EEG from ion channel and circuit level dynamics. Journal of Neuroscience Methods, Methods and Models in Sleep Research: A Tribute to Vincenzo Crunelli, 316, 46–57. 10.1016/j.jneumeth.2018.10.002

Rulkov, N. F. (2002). Modeling of spiking-bursting neural behavior using two-dimensional map. Physical Review E, 65(4), 041922. 10.1103/PhysRevE.65.041922

Rulkov, N. F., & Bazhenov, M. (2008). Oscillations and Synchrony in Large-scale Cortical Network Models. Journal of Biological Physics, 34(3–4), 279–299. 10.1007/s10867-008-9079-y

Rulkov, N. F., Timofeev, I., & Bazhenov, M. (2004). Oscillations in Large-Scale Cortical Networks: Map-Based Model. Journal of Computational Neuroscience, 17(2), 203–223. 10.1023/B:JCNS.0000037683.55688.7e

Steriade, M., Nunez, A., & Amzica, F. (1993). A novel slow (< 1 Hz) oscillation of neocortical neurons in vivo: Depolarizing and hyperpolarizing components. Journal of Neuroscience, 13(8), 3252–3265. 10.1523/JNEUROSCI.13-08-03252.1993

Stevens, C. F. (1993). Quantal release of neurotransmitter and long-term potentiation. Cell, 72, 55–63. 10.1016/S0092-8674(05)80028-5

Van Essen, D. C., Smith, S. M., Barch, D. M., Behrens, T. E. J., Yacoub, E., & Ugurbil, K. (2013). The WU-Minn Human Connectome Project: An overview. *NeuroImage*, Mapping the Connectome, 80, 62–79. 10.1016/j.neuroimage.2013.05.041

Wei, Y., Luo, M., Mai, X., Feng, L., Tang, T., Yang, D., Krishnan, G. P., & Bazhenov, M. (2023). The role of age-related sleep EEG changes in memory decline: Experiments and computational modeling. 2023 45th Annual International Conference of the IEEE Engineering in Medicine & Biology Society (EMBC), 1–4. 10.1109/EMBC40787.2023.10340681

